# Novel ACE2-Independent Carbohydrate-Binding of SARS-CoV-2 Spike Protein to Host Lectins and Lung Microbiota

**DOI:** 10.1101/2020.05.13.092478

**Authors:** Fabrizio Chiodo, Sven C.M. Bruijns, Ernesto Rodriguez, R.J. Eveline Li, Antonio Molinaro, Alba Silipo, Flaviana Di Lorenzo, Dagmar Garcia-Rivera, Yury Valdes-Balbin, Vicente Verez-Bencomo, Yvette van Kooyk

## Abstract

The immediate call for translational research in the field of coronavirus disease (COVID-19) pandemic, needs new and unexplored angles to support and contribute to this important worldwide health problem. The aim of this study is to better understand the pathogenic mechanisms underlying COVID-19, deciphering the carbohydrate-mediated interactions of the SARS-CoV-2 spike protein. We studied the carbohydrate-binding receptors that could be important for viral entry and for immune-modulatory responses, and we studied the interactions of the spike protein with the host lung microbiota. Exploring solid-phase immunoassays, we evaluated the interactions between the SARS-CoV-2 spike protein and a library of 12 different human carbohydrate-binding proteins (C-type lectins and Siglecs) involved in binding, triggering and modulation of innate and adaptive immune-responses. We revealed a specific binding of the SARS-CoV-2 spike protein to the receptors DC-SIGN, MGL, Siglec-9 and Siglec-10 that are all expressed on myeloid immune cells. In addition, because the lung microbiota can promote or modulate viral infection, we studied the interactions between the SARS-CoV-2 spike protein and a library of *Streptococcus pneumoniae* capsular polysaccharides, as well as other bacterial glyco-conjugates. We show specific binding of the spike protein to different *S. pneumoniae* capsular polysaccharides (serotypes 19F and 23F but not to serotype 14). Moreover we demonstrated a specific binding of SARS-CoV-2 spike protein to the lipopolysaccharide from the opportunistic human pathogen *Pseudomonas aeruginosa*, one of the leading cause of acute nosocomial infections and pneumonia. Interestingly, we identified rhamnosylated epitopes as one of the discriminating structures in lung microbiota to bind SARS-CoV-2 spike protein. In conclusion, we revealed novel ACE2-independent carbohydrate-mediated interactions with immune modulating lectins expressed on myeloid cells, as well as host lung microbiota glyco-conjugates. Our results identified new molecular pathways using host lectins and signalling, that may contribute to viral infection and subsequent immune exacerbation. Moreover we identified specific rhamnosylated epitopes in lung microbiota to bind SARS-CoV-2, providing a hypothetical link between the presence of specific lung microbiota and SARS-CoV-2 infection and severity.

## Main Text

Viral infections constitute a serious public health and social problem worldwide, especially during the recent COVID-19 pandemic caused by the betacoronavirus SARS-CoV-2 [1]. This virus utilizes a glycosylated spike protein to bind to the host angiotensin-converting enzyme 2 (ACE2) and TMPRSS2 [2,3]. During the initial and crucial infection steps, different carbohydrate-mediated interactions are playing key roles at the host-pathogen interface, as shown for many viruses such as HIV-1, Influenza viruses, Ebolavirus and SARS-CoV [4,5]. The viral proteins, often highly glycosylated, can be recognized by different carbohydrate-binding proteins from the host, triggering simultaneously immune-responses that can facilitate or inhibit viral entry or dissemination [5]. In contrast to other pathogens and, with rare exceptions, viruses do not have a glycosylation machinery and exploit the host intracellular glycosylation machinery, during the viral intracellular replication. Due to the high evolutionary pressure, the glycosylation profiles of various human-infecting viruses are very different from a classic host-glycosylated protein as described for example for surface proteins of HIV-1, Ebolavirus, MERS, SARS-CoV and SIV [5,6]. The glycans on viral proteins are important for proper folding and for modulating accessibility to host proteases and neutralizing antibodies [4]. Virus have exploited the host-glycosylation machinery as camouflage strategy, hiding hypothetical immunogenic epitopes, as well as using the host carbohydrate binding receptors as entry mechanisms as, for example, described for Ebolavirus [7] and HIV-1 [8]. In addition, low affinity antibodies against some viral glycoproteins have been reported to increase the viral infection [4]. In this scenario, the glycosylation profiles of the spike glycoprotein from SARS-CoV-2 have been recently described [9,10,11]. *N*-glycosylation profile of the SARS-CoV-2 spike protein expressed in human embryonic kidney 293 cells (HEK293) showed a combination of high mannoses and complex-type oligosaccharides, including highly processed sialylated complex-type glycans and fucosylated glycans on different glycosylation sites [9,10,11]. Interestingly, the *N*-glycosylation profile of the SARS-CoV-2 spike protein shows also some difference compared with the SARS-CoV spike protein [11]. In addition, *O*-linked glycan residues have been predicted [12] and described [10]. Apparently three adjacent predicted *O*-linked glycans seem to be unique for the SARS-CoV-2 spike protein.

The host-immune system has an evolutionary conserved class of carbohydrate-binding proteins (lectins) able to sense these different glycan-epitopes, triggering innate responses in a carbohydrate-dependent manner [13]. In this frame, the C-type lectins dendritic cell-specific intercellular adhesion molecule-3-grabbing non-integrin (DC-SIGN) and Langerin have been described as well-known receptor on dendritic cells used by many viruses as entry mechanism such as HIV-1 [14,15] and Ebolavirus [7]. The macrophage galactose binding lectin (MGL) and the mannose receptor (MR) play a role in Influenza A virus infection [16], and sialic acid-binding immunoglobulin-type lectins (Siglecs)-1 has been involved in HIV-1 *trans* infection and Ebolavirus uptake [17,18].

In the context of SARS-CoV-2 infection, we investigated the interactions between the glycosylated viral spike protein expressed in HEK293 cells and the host immune lectins via solid-phase immunoassays. We screened a library of different human C-type lectins expressed on myeloid cells such as the antigen-presenting cells like dendritic cells and macrophages (DC-SIGN, Langerin, MGL, MR, Dectin-1 and the macrophage inducible calcium dependent lectin receptor Mincle) or soluble (MBL), to bind to the SARS-CoV-2 spike protein used as coating agent in ELISA experiments. **Figure 1** clearly shows binding of the SARS-CoV-2 spike protein to human lectin receptors DC-SIGN and MGL. The binding to DC-SIGN (but not to Langerin, MR and MBL) suggests the interactions with specific fucosylated glycans present at N343 (core α1,6 fucose, or with LeX type of structures) [9] or a particular conformation of the high-mannoses able to discriminate between DC-SIGN and Langerin. The binding of the SARS-CoV-2 spike protein to MGL fits with the presence of *O*-glycans at Thr323 or Ser325 on the spike protein [10], as the specificity of MGL has been described for α or β Gal/GalNAc containing glyco-conjugates [19]. Surprisingly both MGL and DC-SIGN are receptors found on tolerogenic macrophages/dendritic cells and their carbohydrate-binding have been shown to hijack myeloid function keeping macrophages immune-silent by the induction of anti-inflammatory cytokines and induction of T cell apoptosis/exhaustion [20]. Although the oligomannosides-content of the spike protein has been reported to be around 30% and Man_5_GlcNAc_2_ has been described to be the predominant oligomannose-type glycan (N234 and N709) [9], we did not see binding to Langerin, MR and MBL being *bona fide* lectins able to recognize mannosides. Despite the fact that the MR mannose-binding evolved separately from the other C-type lectins, MBL and MR have close similarity in their mannose-binding profiles [21]. However, these human lectins were able to bind the positive controls of our experiment, illustrating sensitivity of the assay for these lectins to bind their carbohydrate structures (see Figure S1). The specific binding to DC-SIGN but not to Langerin has been shown also for Marburg virus (MARV) [22], while viruses like HIV-1 binds both receptors [15]. The fact that SARS-CoV-2 spike protein binds DC-SIGN but not Langerin nor MR or MBL, may indicate that the high-mannoses are not accessible or that there is a preferential specificity for the different fucosylated residues in the context of core fucosylation or terminal Lewis antigens on the spike protein. This is line with our finding that both lectins do not bind to the spike protein confirming poor accessibility of the spike protein high-mannoses to the immune system. As expected, the lectin Mincle and Dectin-1, recognising mostly mycobacterial glycolipids and beta-glucans respectively, were used as control and did not bind to the spike protein. As full figure with all the positive controls is shown in **Figure S1**.

**Figure 1:**
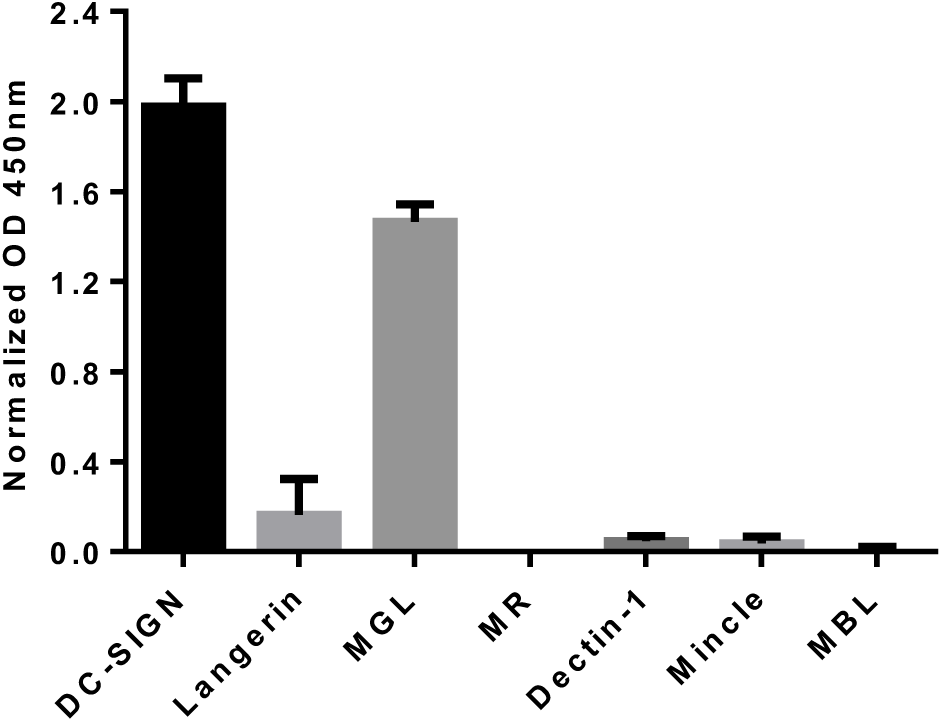
Binding of SARS-CoV-2 spike protein to human C-type lectins. Human C-type lectins ELISA results: wells were coated with the viral spike protein and the binding to different C-type lectins was evaluated in a calcium-containing buffer. The experiment has been performed three times in duplicate with similar results, and data were normalized over signal from BSA-coated wells. Error bars indicate standard deviations. OD: Optical density.

Since the SARS-CoV-2 spike protein has been described to express highly processed sialylated complex-glycans at N165, N282, N801, N1074 and N1098 [10] we investigated whether these glycans could be a signature for the sialic-acid binding lectins (Siglecs), a family of carbohydrate receptors with immune-receptor tyrosine-based inhibition motif (ITIM) and known immune inhibitory signaling properties [23]. Using our solid-phase immunoassay screening-system, a library of different Siglecs (Siglec-3, Siglec-5, Siglec-7, Siglec-9 and Siglec-10) was explored for binding activity to SARS-CoV-2 spike protein. Siglecs can be expressed on almost all immune cells, including B and T cells, monocytes, eosinophils, and macrophages [23]. Although Siglecs are *bona fide* Sialic-acids binding proteins, the literature is not consistent in assigning a specific sialylated pattern to each Siglec. Notwithstanding this, human Siglec-3 and Siglec-10 have a prevalence to interact with α2,6 linked sialic acids while Siglec-9 has a prevalence to interact with α2,3 linked sialic acids (in the context, in most of the cases, of fucosylated or sulfated oligosaccharides). Siglec-3 is highly expressed on macrophages and monocytes; Siglec-9 on monocytes, neutrophils, conventional dendritic cells, NK and T cells; Siglec-10 on B cells, monocytes and eosinophils [24]. Different Siglecs have been implicated in T cell-mediated tolerance, alter B cell function and the inhibitory Siglec-9 and Siglec-10 were upregulated on tumor-infiltrating CD4+ and CD8+ T cells contributing to exhaustion of tumor T cell responses [23].

As shown in **Figure 2** we observed a specific interaction of the SARS-CoV-2 spike protein with Siglec-3, Siglec-9 and Siglec-10. Our results clearly suggest the presence of α2,6 and α2,3 linked sialic acids on the viral spike protein, as well as the roles of immunomodulatory Siglecs during SARS-CoV-2 infection, probably by altering and modulating the B-cells, neutrophil, eosinophils, monocytes and macrophages function. As full figure with the positive controls is shown in **Figure S2**.

**Figure 2:**
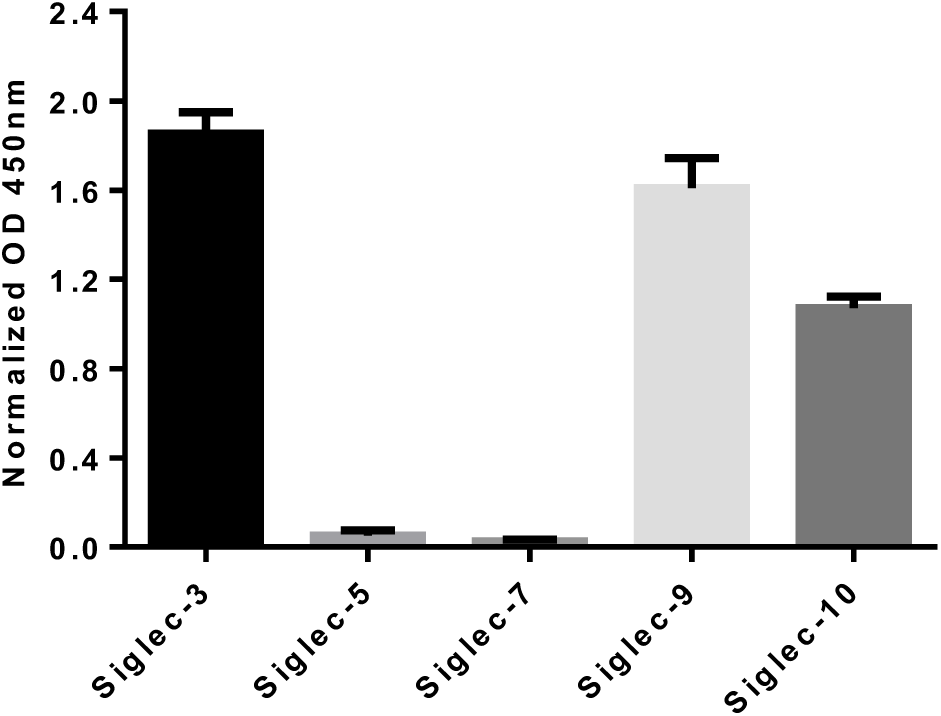
Binding of SARS-CoV-2 spike protein to human Siglecs. Human Siglecs ELISA results: wells were coated with the viral spike protein and the binding to different Siglces was evaluated. The experiment has been performed three times in duplicate with similar results, and data were normalized over signal from blocking buffer-coated wells. Error bars indicate standard deviations. OD: Optical density.

These data illustrate that the heterogeneous glycan-repertoire expressed on the SARS-CoV-2 spike protein modulate the binding to different carbohydrate-binding receptors expressed on host immune cells. In particular we speculate that the spike protein could target a repertoire of immune inhibiting glycan binding receptors, such as DC-SIGN, MGL, Siglec-3, Siglec-9 and Siglec-10. This also illustrates the enormous potential of SARS-CoV-2 to modulate the host immune system on multiple levels.

We next explored a different angle of carbohydrate-mediated interactions of SARS-CoV-2 studying the interactions with lung microbiota components that may contribute to the disease severity. Various viruses such as Influenza viruses, HIV-1, coronavirus OC43 and MERS, have been described to interact with the host sialic-acids containing structures [25, 26] or to interact with sulfated glycosaminoglycans, suggesting a carbohydrate-recognition activity for different viral proteins [27]. In this context, recently the glycosaminoglycan heparin has been described to interact with the SARS-CoV-2 spike protein inducing conformation changes [28] and heparan sulfate showed binding to the spike protein in a length dependent manner [29]. Starting from these evidences, we chose to study the interaction of the SARS-CoV-2 spike protein with different glyco-conjugates from the lung microbiota. The mechanism underlying the interactions between the lung microbiota and viruses are currently not fully understood. It is known that the lung resident microbiota modulates viral infections yielding beneficial outcomes, conferring protection against viral infection, or harmful outcomes since virus can use the resident bacteria to spread in the host by direct or indirect mechanisms [30,31,32,33]. Moreover, viruses can perturb the integrity of the commensal microbiota (modified by viral infections, age, and smoke), that in turn influences virus infectivity and morbidity [31]. Looking closely to all the bacteria occupying the lung and upper-respiratory microbiota, they are covered by an extremely complex library of glyco-conjugates mostly in the form of bacterial capsular-polysaccharides (CPS) and lipopolysaccharides (LPS). This outer layer may contribute to the intimate interactions between SARS-CoV-2 and the lung microbiota, and could help not only in understanding mechanistically the viral entry, but also in supporting therapies and prognosis/diagnosis markers. From a clinical perspective, few data suggest that the microbiota composition of the upper-respiratory tract during acute respiratory syncytial virus infection differs from that of healthy controls [34]. During the COVID-19 pandemic, as well as other past viral outbreaks, respiratory viral infections are associated with co-infections with *Streptococcus pneumoniae* (Sp). As recently reported, 50% of COVID-19 patients who have died, also showed secondary bacterial infections, and fungal co-infections [35].

Thus, as a proof-of-concept, we designed a solid-phase immunoassay coating the plates with a library of seven structurally different CPS from *S. pneumoniae* (the seven most abundant serotypes worldwide as well as the ones included in pneumococcal conjugate-vaccines, PCV: Sp1, Sp5, Sp06B, Sp14, Sp18C, Sp19F and Sp23F) and determined the binding of SARS-CoV-2 spike protein as Fc-construct in our assay to quantify potential interactions. Surprisingly, when analyzing the binding of the spike protein to the structurally different Sp CPS with our assay, we observed a strong binding of the spike protein to Sp19F and Sp23F (**Figure 3**). The spike protein was able to strongly recognize the CPS from Sp19F and Sp23F, while a somewhat weaker interaction was observed with Sp18C and Sp06B. The CPS from Sp1 and Sp5 poorly bound SARS-CoV-2 spike protein, while the CSP from Sp14 was not recognized. Looking for a structural rationale, Sp19F, Sp23F and Sp18C, Sp06B have in common the presence of the following glycan epitope: Glc (or Gal) 1,2 (1,3 or 1,4) linked to Rham (where Rha stands for Rhamnose, Glc for Glucose and Gal for Galactose). Interestingly, Sp1, Sp5 and Sp14 have no rhamnose in their structures. Rhamnose is a deoxy sugar not present in humans but highly abundant for example in bacteria and mycobacteria. Bacterial rhamnosylated glyco-conjugates have evolutionary conserved patterns that co-evolved with the host immune system, triggering for example, in mammals a conserved and abundant class on naturally-acquired anti-rhamnose antibodies [36]. Despite the fact that we see differential binding of the SARS-CoV-2 spike protein to these different capsules, we are sure that an equal amounts is coated in the plates: increase of the concentrations of the different a CPS showed no improvement of the binding, and different monoclonal antibodies against the reported CPS have been tested in these experimental conditions showed equal coating of the different CPS.

**Figure 3:**
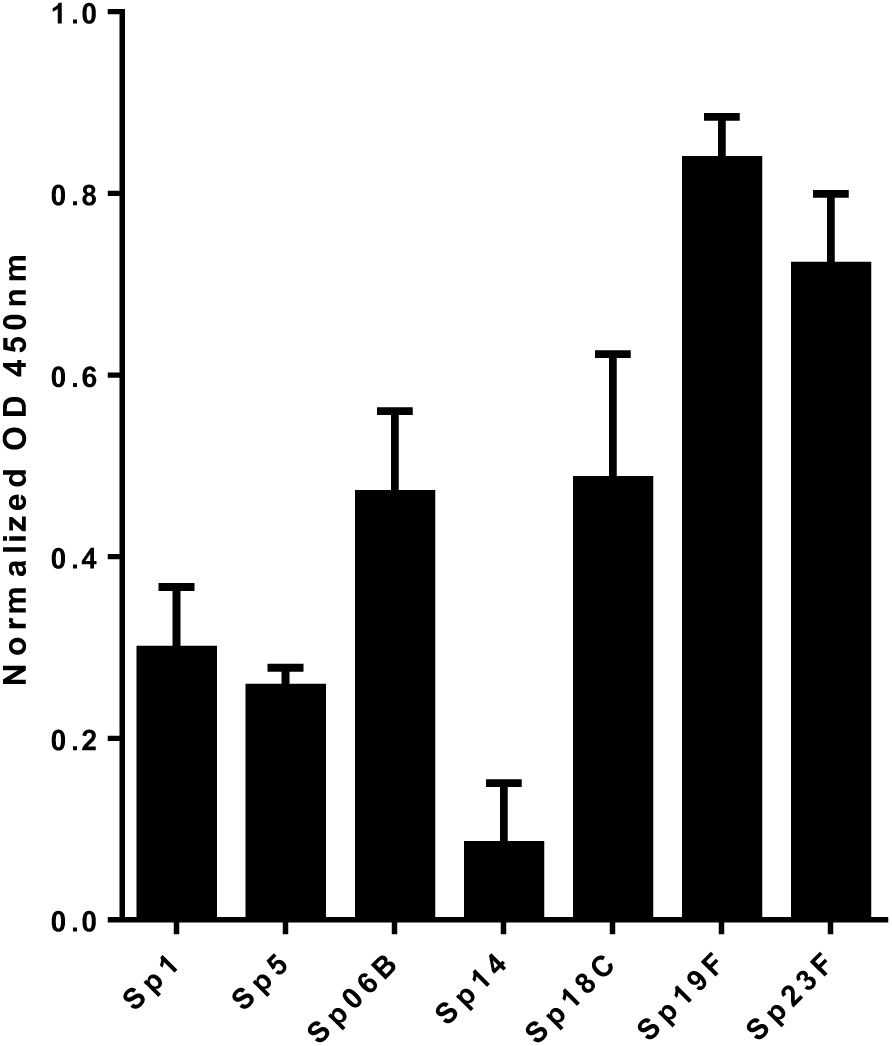
Binding of SARS-CoV-2 spike protein to different *S. pneumoniae* CPS. Wells were coated with different *S. pneumoniae* CPS and the binding to the viral spike protein was evaluated exploiting the mouse-Fc tag on the recombinant chimera spike protein. The experiment has been performed three times in duplicate with similar results, and data were normalized over signal from blocking buffer-coated wells. Error bars indicate standard deviations. OD: Optical density.

Following the observation of the binding of the SARS-CoV-2 spike protein to different Sp CPS, we aimed at studying also the interactions between other lung microbiota glyco-conjugates. We designed another ELISA experiment to prove the spike protein binding to different LPS expressed as surface antigens from different Gram-negative bacteria. A small library of different LPS (*Pseudomonas aeruginosa, Salmonella typhimurium* and *Shigella flexneri*) was tested in an ELISA assay against the spike protein. *P. aeruginosa* is a typical and widespread human opportunistic pathogen, it can cause severe and often deadly lung infections, particularly in immunocompromised patients, and increasingly resistant to a large number of antibiotics [37]. We chose to compare *P. aeruginosa* to two bacteria not interesting the lungs, namely S. *typhimurium* [38] and *S. flexneri* [39], enteric pathogens both capable of invading and disseminating the intestinal epithelium. The specific interaction between the spike protein and the LPS from *P. aeruginosa*, when compared to the LPS from the gut-associated pathogens *S. typhimurium* and *S. flexneri*, is indeed clear and showed in **Figure 4**. The same structural considerations mentioned for the Sp CPS might be here done, that is the LPS from the intestinal pathogens here tested (*Salmonella* and *Shigella*) do not express rhamnosylated-epitopes, while the *P. aeruginosa* strain here analysed. In addition to be an opportunistic pathogen able to occupy the lung ecosystem, *P. aeruginosa* expresses a LPS containing in its core-antigen the Glc 1,2-Rha epitope [40]. Also in this case, we proved with different parallel experiments (lectins binding) that the three tested LPS are coating the ELISA wells in a similar way.

**Figure 4:**
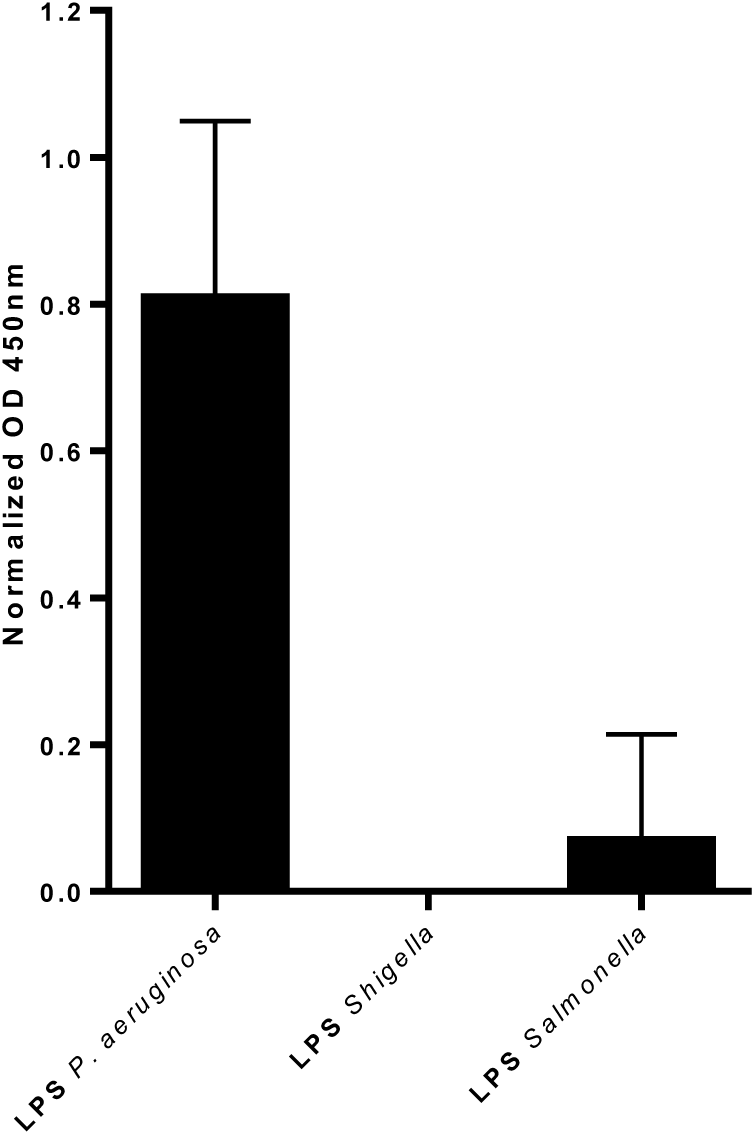
Binding of SARS-CoV-2 spike protein to different LPS. Wells were coated with different LPS and the binding to the viral spike protein was evaluated exploiting the mouse-Fc tag on the recombinant chimera spike protein. The experiment has been performed three times in duplicate with similar results, and data were normalized over signal from blocking buffer-coated wells. Error bars indicate standard deviations. OD: Optical density.

All together our results on bacterial glyco-conjugates suggest, in the context of SARS-CoV-2 infection, that some bacteria (and serotypes) in the lung, that harbor a rhamnosylated-epitopes, could facilitate the binding of the virus to the epithelial cells and its subsequent dissemination. These observations could trigger the design of new therapeutic approaches for the bacterial co-infections in COVID-19 patients. These unreported results indicate that lung microbiota CPS and LPS, can play a role in accelerating the viral infection. In addition, due to the very high prevalence of bacterial pneumonia in COVID-19 patients, these results could also help in evaluating the impact of PCV vaccination on COVID-19 patients (as already suggested of other viral infections [41]), as well as to identify specific Sp serotypes able to modulate the viral infection.

In conclusion, we identified different carbohydrate-mediated interactions of the SARS-CoV-2 spike protein. These interactions could impact and drive the infection severity and the corresponding uncontrolled immune responses and inflammation processes [42].

Here we proved that the glycan-heterogeneity on the SARS-CoV-2 spike protein is the cause of the multiple interactions with the host lectins. We showed that the spike protein is recognized specifically by C-type lectins (DC-SIGN and MGL) and by different Siglecs (Siglec-3, Siglec-9 and Siglec-10). Interestingly this connects to new, ACE2 independent, infection pathways, with the hypothetical immune suppression of monocytes, macrophages, dendritic cells and B cells. The binding to DC-SIGN, MGL, Siglec-9 and Siglec-10, indicates the presence and accessibility of the immune system, to fucosylated and sialylated structures as well as the clear presence of *O*-glycans on the spike glycoprotein.

We also proved a totally unexplored behavior for viral proteins, showing the specific binding of the SARS-CoV-2 to different glyco-conjugates expressed by the bacteria able to colonize the lungs. These results may have implications to control and understand the bacterial co-infections, to have a better understanding on the disease severity and to add risk factors in COVID-19 patients.

Together these data aim at adding different carbohydrate-mediated interactions in the complex and holistic picture of the SARS-CoV-2 infection and its implications.

## Acknowledgements

This work was Financial supported by Immunoshape (MSCA-ITN-2014-ETN No 642870) to E.R., European Research Council (ERC-339977-Glycotreat) to Y.K., European Research Councel-MATCH-862113 to E.L., and by NWO Spinoza award to F.C. and E.R.

## Experimental: C-type lectins ELISA

a solution of 50 µL of SARS-CoV-2 spike protein (2019-nCoV Spike RBD-His Recombinant Protein, Cat: 40592-V08H, expressed in HEK293 cells, purchased from Sinobiological) at 5 µg/mL, in PBS (10mM, pH=7.4), were used to coat the Nunc MaxiSorp plate 2h at 37°C. After discarding and washing (2×150µL) with calcium and magnesium-containing buffer TSM [20 mM tris(hydroxymethyl)aminomethane (Tris)-HCl, pH 8.0; 150 mM NaCl; 1 mM CaCl_2_; 2 mM MgCl_2_], the wells were blocked with 100 µL of 1% BSA (Sigma-Aldrich, lyophilized powder, ≥96%, agarose gel electrophoresis) in TSM at 37°C for 30 min. The blocking solution was discarded and 50µL of the different C-type-lectins human-Fc at 1 µg/mL in assay buffer (TSM, 0.5% BSA) were added to the wells. After 1h at room temperature, the wells were washed with TSM (2×150µL) and then 100 µL of anti-human horseradish peroxidase (0.3 µg/mL, Goat anti-Human IgG-HRP from JacksonImmuno) were added. After 30 min at room temperature, the wells were washed with TSM (2×150µL). Finally, 100 µL of substrate solution (3,3′,5,5′-Tetramethylbenzidine, TMB, in citric/acetate buffer, pH=4, and H_2_O_2_) were added and after max 15 min incubation at room temperature the reaction was stopped with 50 µL of H_2_SO_4_ (0.8 M) and the optical density (OD) was measured at 450 nm in an ELISA reader. The experiment was performed three times with similar results, in duplicate and data were normalized over the signal at 450 nm from the BSA-containing wells. Polyacrylamide polymers (PAA), functionalized with different glycans were purchased from Lectinity, MW approx. 20 KDa, Carbohydrate content around 20% mol. Galβ1-4(Fucα1-3)GlcNAcβ-OCH_2_CH_2_CH_2_NH_2_ 0044-PA (PAA-LeX, positive control for DC-SIGN), and GalNAcα-OCH_2_CH_2_CH_2_NH_2_ 0030-PA (PAA-Tn, positive control for MGL). We used a concentration of 40μg/mL PAA-glyco-conjugates to coat the ELISA wells. Laminarin from *Laminaria digitata* (positive control for Dectin-1) and Mannan from *Saccharomyces cerevisiae* (positive control for DC-SIGN, Langerin, MR and MBL) were purchased from Sigma-Aldrich. Trehalose-6,6-dibehenate (TDB, positive control for Mincle) was purchased from Invivogen. For these three molecules, we used a concentration of 40μg/mL to coat the ELISA wells.

## Siglecs ELISA

a solution of 50 µL of SARS-CoV-2 spike protein (2019-nCoV Spike RBD-His Recombinant Protein, Cat: 40592-V08H, expressed in HEK293 cells, purchased from Sinobiological) at 5 µg/mL, in PBS (10mM, pH=7.4), were used to coat the Nunc MaxiSorp plate 2h at 37°C. After discarding and washing (2×150µL) with Hanks’ Balanced Salt solution (Gibco™ HBSS) the wells were blocked with 200 µL of carbo-free blocking solution (Vector Laboratories, catalog No.NC9977573) at 37°C for 30 min. The blocking solution was discarded and 50µL of the different human Siglecs-Fc constructs (recombinant human chimera purchased from R&D Systems Netherlands) at 5 µg/mL (for Siglec-9 100ng/mL) in assay buffer (carbo-free solution) were added to the wells. After 2h at room temperature, under gentle shaking (150 rot/min), the wells were washed with HBSS (2×150µL) and then 100 µL of anti-human horseradish peroxidase (0.3 µg/mL, Goat anti-Human IgG-HRP from JacksonImmuno) were added. After 30 min at room temperature, the wells were washed with HBSS (2×150µL). Finally, 100 µL of substrate solution (3,3′,5,5′-Tetramethylbenzidine, TMB, in citric/acetate buffer, pH=4, and H_2_O_2_) were added and after max 15 min incubation at room temperature the reaction was stopped with 50 µL of H_2_SO_4_ (0.8 M) and the optical density (OD) was measured at 450 nm in an ELISA reader. The experiment was performed three times with similar results, in duplicate and data were normalized over the signal at 450 nm from the carbo-free blocking solution-containing wells. Polyacrylamide polymers (PAA), functionalized with different glycans were purchased from Lectinity, MW approx. 20 KDa, Carbohydrate content around 20% mol. Neu5Acα6’Lac-C_2_-PAA 0063a-PA (PAA-Sia 2,6), Neu5Acα3’Lac-Gly-PAA 0060-PA (PAA-Sia 2,3), and Galβ-OCH_2_CH_2_CH_2_NH_2_ 0024-PA (PAA-Gal). We used a concentration of 40μg/mL PAA-glyco-conjugates to coat the ELISA wells.

## Spike protein bacterial glyco-conjugates ELISA

a solution of 50 µL of different bacterial glyco-conjugates (*S. pneumoniae* CPS were provided by the Finlay Vaccine Institute, Cuba. The LPS from *Pseudomonas aeruginosa* serotype 10, *Shigella flexneri* strain M90T, *Salmonella typhimurium* SH2201, were provided by the Department of Chemical Sciences, University of Naples Federico II, Italy) at 40 µg/mL (CPS) or 20 µg/mL (LPS), in PBS (10mM, pH=7.4), were used to coat the Nunc MaxiSorp plate 2h at 37°C. After discarding and washing (2×150µL) with PBS (10mM, pH=7.4), the wells were blocked with 100 µL of carbo-free blocking solution (Vector Laboratories, catalog No.NC9977573) at 37°C for 30 min. The blocking solution was discarded and 50µL of SARS-CoV-2 spike protein mice-Fc chimera (2019-nCoV Spike Protein RBD, Fc Tag, purchased from Sinobiological) at 2 µg/mL in carbo-free buffer were added to the wells. After 1h at room temperature, the wells were washed with PBS (2×150µL) and then 100 µL of anti-mouse horseradish peroxidase (1 µg/mL, Goat anti-mouse IgG-HRP from JacksonImmuno) were added. After 30 min at room temperature, the wells were washed with PBS (2×150µL). Finally, 100 µL of substrate solution (3,3′,5,5′-Tetramethylbenzidine, TMB, in citric/acetate buffer, pH=4, and H_2_O_2_) were added and after max 15 min incubation at room temperature the reaction was stopped with 50 µL of H_2_SO_4_ (0.8 M) and the optical density (OD) was measured at 450 nm in an ELISA reader. The experiment was performed three times with similar results, in duplicate and data were normalized over the signal at 450 nm from the carbo-free blocking solution-containing wells.

