## Supplementary Material for "Novel ACE2-Independent Carbohydrate-Binding of SARS-CoV-2 Spike Protein to Host Lectins and Lung Microbiota"

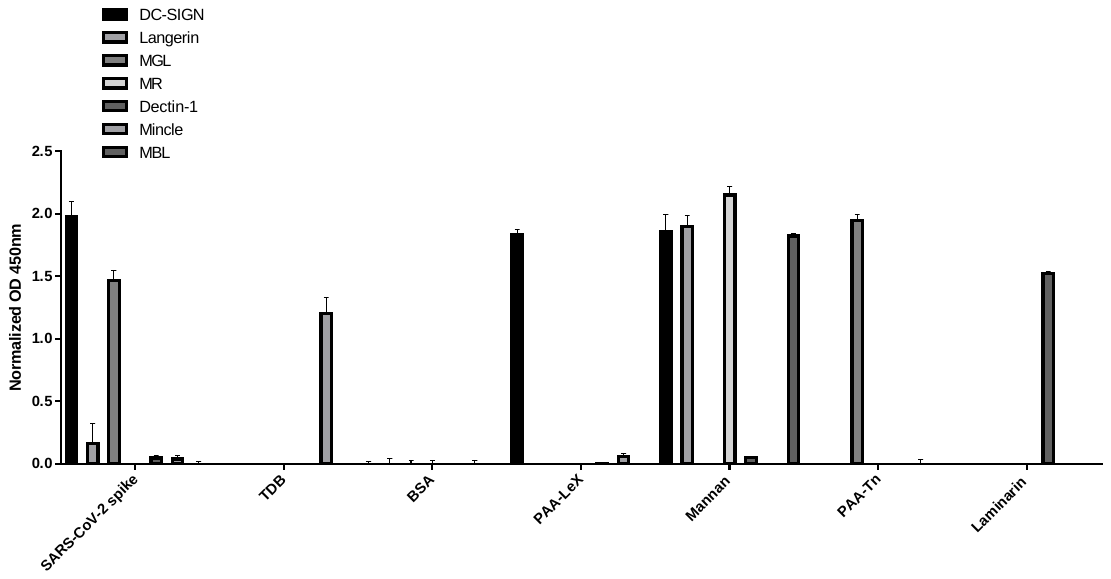

**Figure S1:** Binding of SARS-CoV-2 spike protein to human C-type lectins. Human C-type lectins ELISA results: wells were coated with the viral spike protein and the binding to different C-type lectins was evaluated in a calcium-containing buffer. The experiment has been performed three times in duplicate with similar results, and data were normalized over signal from BSA-coated wells. Error bars indicate standard deviations. OD: Optical density.

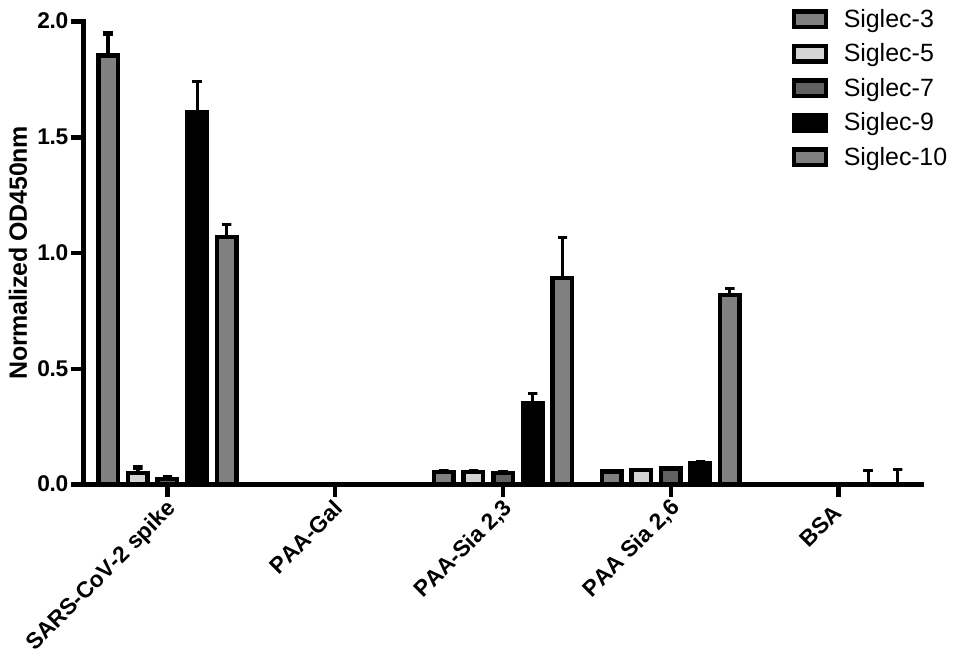

**Figure S2:** Binding of SARS-CoV-2 spike protein to human Siglecs. Human Siglecs ELISA results: wells were coated with the viral spike protein and the binding to different Siglecs was evaluated. The experiment has been performed three times in duplicate with similar results, and data were normalized over signal from blocking buffer-coated wells. Error bars indicate standard deviations. OD: Optical density.

The high positive binding of the viral spike protein to the different lectins in our experiments, was comparable to all the positive controls used. We are aware that ELISA are not quantitative experiments especially comparing different macromolecules (spike protein, Mannan etc), but having the positive results in the same OD range, provides a solid interpretation of our data.
